# Loss of the cleaved-protamine 2 domain leads to incomplete histone-to-protamine exchange and infertility in mice

**DOI:** 10.1101/2021.09.29.462440

**Authors:** Lena Arévalo, Gina Esther Merges, Simon Schneider, Franka Enow Oben, Isabelle Neumann, Hubert Schorle

## Abstract

Protamines are unique sperm-specific proteins that package and protect paternal chromatin until fertilization. A subset of mammalian species expresses two protamines (PRM1 and PRM2), while in others PRM1 is sufficient for sperm chromatin packaging. Alterations of the species-specific ratio between PRM1 and PRM2 are associated with infertility. Unlike PRM1, PRM2 is generated as a precursor protein consisting of a highly conserved N-terminal domain, termed cleaved PRM2 (cP2), which is consecutively trimmed off during chromatin condensation. The carboxyterminal part, called mature PRM2 (mP2), interacts with DNA and together with PRM1, mediates chromatin-hypercondensation. The removal of the cP2 domain is believed to be imperative for proper chromatin condensation, yet, the role of cP2 is not yet understood. We generated mice lacking the cP2 domain while the mP2 is still expressed. We show that the cP2 domain is indispensable for complete sperm chromatin protamination and male mouse fertility. cP2 deficient sperm show incomplete PRM2 incorporation, resulting in a severely altered protamine ratio, retention of transition proteins and aberrant retention of the testis specific histone variant H2A.L.2. During epididymal transit, cP2 deficient sperm seem to undergo ROS mediated degradation leading to complete DNA fragmentation. The cP2 domain therefore seems to be a key aspect in the complex crosstalk between histones, transition proteins and protamines during sperm chromatin condensation. Overall, we present the first step towards understanding the role of the cP2 domain in paternal chromatin packaging and open up avenues for further research.

## Introduction

Chromatin structure and dynamics in the sperm nucleus are as unique as the sperm cell itself, and are of major importance to sperm function, fertilizing ability, and embryo survival. Paternal DNA is particularly vulnerable to damage, especially to oxidative stress during epididymal sperm maturation and migration, leading to a higher requirement for protection (Chen et al. 2002). At the same time, the size and shape of the sperm cell nucleus needs to be optimized for efficient sperm movement through the female reproductive tract (Tourmente et al. 2011). This is achieved by the complete reorganization of paternal chromatin from nucleo-histone to nucleo-protamine during the final steps of spermatogenesis (Balhorn 2007, Rathke et al. 2014).

Even though many studies have demonstrated that the correct execution of this transition is imperative for male fertility and embryo survival (Aoki and Carrell 2003), surprisingly little is known about the process itself. Recent studies have only just started to unravel its molecular basis (Schneider et al. 2016, Barral et al. 2017, Hada et al. 2017). Barral et al. (2017) were able to show that the testis specific histone variant H2A.L.2 together with TH2B and transition proteins (TNP1 and TNP2) mediates structural changes in chromatin allowing protamines to bind DNA. Protamines are small, arginine-rich proteins. Their high arginine content allows them to bind DNA with high affinity and to shield the charges of the DNA backbone more efficiently than histones (Balhorn 1989, Tanaka and Baba 2005). Two types of protamines have been identified in mammals: protamine 1 *(PRM1,* PRM1) and protamine 2 *(PRM2,* PRM2). While PRM1 is a major sperm protamine found across mammals, PRM2 is only detected in the sperm of primates, most rodents, and a subset of other placental mammals (Chauviere et al. 1992, Retief and Dixson 1993). The coding regions of *PRM1* and *PRM2* are tightly clustered and map to a small section of chromosome 16. It is highly likely that *PRM2* is the result of a *PRM1* duplication event (Krawetz and Dixon 1988, Lüke et al. 2011).

Unlike PRM1, PRM2 is transcribed as a precursor. The N-terminal region of the translated PRM2, termed cleaved-PRM2 (cP2), is successively cleaved over several days while chromatin condensation is taking place. After this, only mature-PRM2 (mP2) remains bound to the completely condensed DNA (Retief et al. 1993, Balhorn 2007, Yelick et al. 1987, Oliva and Dixon 1991). Perturbations of PRM2 processing has been shown to lead to decreased DNA integrity and sperm dysfunction (deYebra et al. 1998, Torregrosa et al. 2006). In previous comparative evolutionary studies, it was shown that the cP2 coding sequence is conserved in both primates and rodents. mP2, however, evolves under less selective constraint (Lüke et al. 2011,2016). Changes in coding sequences of cP2 were associated with differences in sperm head size in mouse species. This association was specific to the cP2 domain and not found for mP2 (Lüke et al. 2014a). A potential reason for this pattern is that changes in cP2 are selected against conserving a crucial function for reproduction, while mP2 is free to evolve under less constraint due to its proposed functional redundancy to PRM1 (Lüke et al. 2011,2016).

Proper PRM2 cleaving therefore seems to be crucial for successful reproduction, yet, the function of the cP2 domain and PRM2 processing are unknown to date. Establishment and analysis of PRM2 deficient mice revealed that Prm2^-/-^ males were infertile, while Prm2^+/-^ males remained fertile (Schneider et al. 2016). Of note, mice deficient for TNP1, TNP2 and H2A.L.2 show incomplete PRM2 processing (PRM2 precursor detected in mature sperm nuclei) (Yu et al. 2000, Zhao et al. 2001,2004, Barral et al. 2017).

Given that cP2 cleaving is taking place during DNA condensation in late spermiogenesis, the strong evolutionary conservation of this domain and the effect of incomplete PRM2 processing on sperm function and fertility, cP2 is likely to play an important role in the correct execution of chromatin condensation. We therefore studied the involvement of cP2 in chromatin condensation during spermiogenesis by generating and analyzing a mouse line bearing a deletion of cP2, while maintaining mP2 expression. We analyzed fertility, testis and sperm morphology, chromatin integrity and nuclear protein content of these mice and revealed that Prm2^Δc/+^ mice are already infertile and that cP2 seems necessary for complete protamination of sperm chromatin.

## Material and Methods

### Animals

For the generation of mouse lines using CRISPR/Cas9 the F1 generation of mouse strains C57Bl6 and DBA2 (B6D2F1) were used. Founder animals were backcrossed to C57Bl6. Mice were maintained under standard laboratory conditions in environmentally controlled rooms (20–24°C) on a 12L:12D photoperiod with nesting material and ad libitum food and water. All animal experiments were conducted according to German law of animal protection and in agreement with the approval of the local institutional animal care committees (Landesamt für Natur, Umwelt und Verbraucherschutz, North Rhine-Westphalia, Germany, AZ81-02.04.2018.A369).

### Gene-edited mouse lines

Guide RNA (gRNA) sequence pre-selection was performed using the algorithm published by Hsu et al. (2013). Two gRNA sequences were selected based on quality scores in each case maximizing specificity and minimizing off-target action (score > 50) (Table S1). The designed guide sequences were ordered as crisprRNA sequences (crRNA) (IDT, Leuven, Belgium) and annealed to tracrRNA (IDT) by incubating 5min at 95°C for a final concentration of 50mM of gRNA (crRNA+tracrRNA). The target site of the designed gRNAs is shown in figure S1A. The repair template used for homology directed repair (HDR), here single-stranded oligodeoxynucleotides (ssODNs) (IDT) (Yoshimi et al. 2014, 2016), are shown in figure S1B. Ribonucleoprotein (RNP) complexes were assembled immediately prior to delivery by incubation of 4pmol/μl Cas9 (IDT) protein, 4pmol/μl of each gRNA and 10pmol/μl ssODN in Opti-MEM medium (Thermo Fisher Scientific, Waltham, USA) for 10min at room temperature.

To generate gene-edited founder animals, B6D2F1 females were hormonally superovulated and mated as described (Schneider et al. 2016). Oocytes were isolated from the oviducts, washed and transferred into droplets of Opti-MEM medium containing the previously prepared RNP complex and electroporated using a BioRad Gene Pulser (BioRad, Feldkirchen, Germany) (two 3ms square wave pulses at 30V with an 100ms interval). Oocytes were recovered and washed 5x in M2 medium (Merck Millipore, Darmstadt, Germany) followed by 3 washes in KSOM (Merck Millipore) medium droplets. Oocytes were then incubated in KSOM medium covered in paraffin oil overnight at 37°C. Developing 2-cell stage embryos were then transferred into the oviducts of pseudo-pregnant foster mice. Offspring was genotyped (primers, see Fig. 1A, Table S1) and positive founder animals backcrossed to C57Bl/6J for at least 3 generations before analysis. Male mice between 10 and 13 weeks of age were used for analysis.

**Figure 1.**
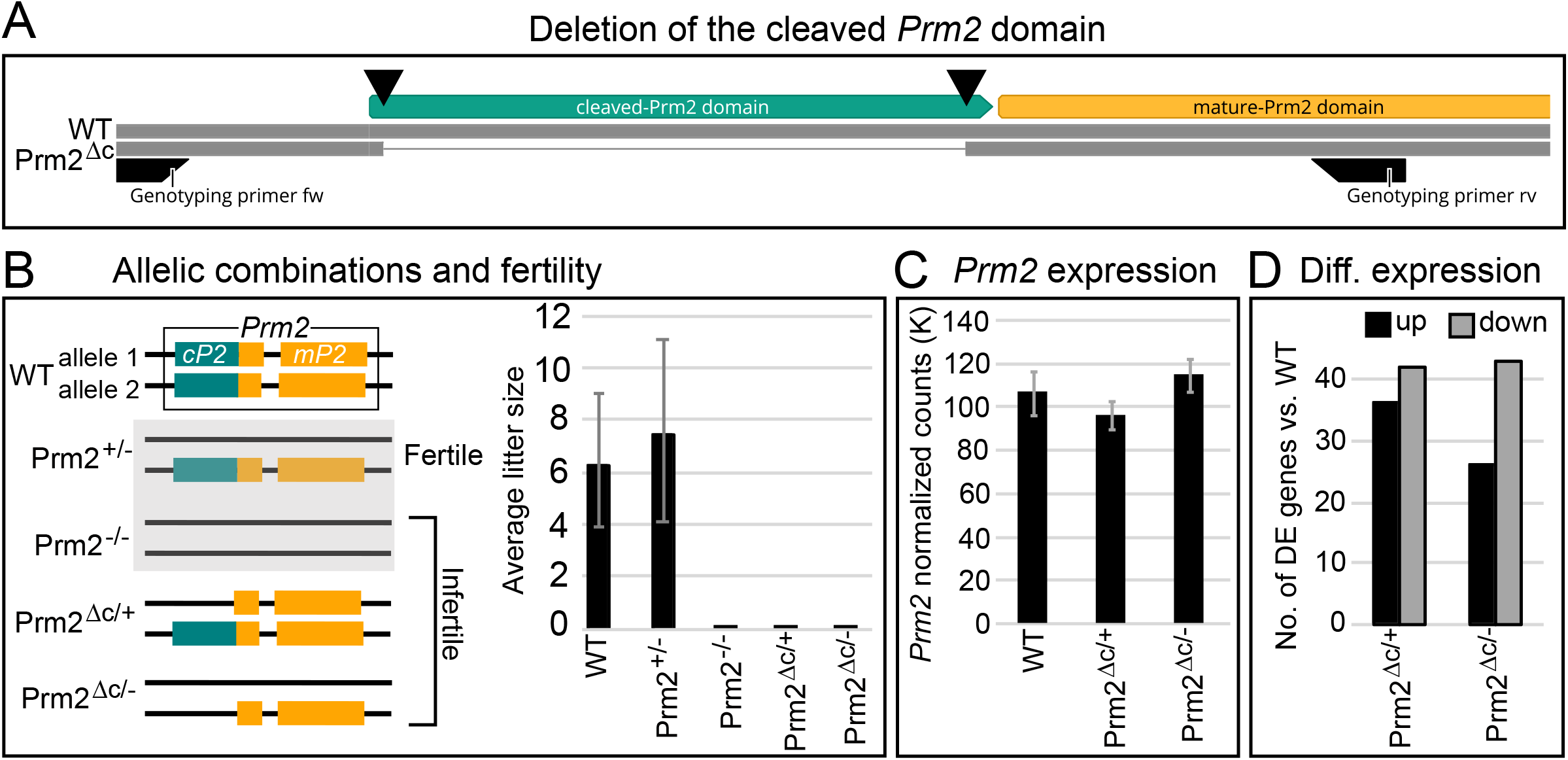
Gene editing, fertility and expression. A) Schematic representation of the generation of cP2 deletion. Double strand breaks induced by Cas9 indicated by black triangles. B) Schematic overview of analyzed genotypes and fertility. Prm2^+/-^ and Prm2^-/-^ (Schneider et al. 2016) were included as a comparison. Barplot of average litter size for WT, Prm2^+/-^, Prm2^-/-^, Prm2^Δc/+^ and Prm2^Δc/-^, n=5 for each genotype. C) Barplot showing average DESeq2 normalized read counts of *Prm2* for WT, Prm2^Δc/+^ and Prm2^Δc/-^. D) Barplot showing comparison between number of differentially higher and lower expressed genes for Prm2^Δc/+^ and Prm2^Δc/-^ compared to wildtype.

### Fertility analysis

Five males per genotype were mated with C57Bl/6J females 1:2 and females were checked daily for the presence of a vaginal plug until at least 5 plugs per male were observed. Pregnancy rate and litter size were noted.

### Sampling and mature sperm isolation

The testes were dissected and weighed. For paraffin sectioning, testes and epididymides were fixed in either Bouin’s solution for histology or in 4% Paraformaldehyde solution for immunohistochemistry (IHC). For tubule preparations testes were transferred to PBS and tubules dissected as described in Kotaja et al. (2004). Briefly, tubules were separated and elongating and condensed spermatid containing sections were identified through their light absorption pattern using a dissection microscope. Tubule sections were squashed on a slide, frozen in liquid nitrogen for 20 seconds, fixed in 90% ethanol for 5 minutes and air dried.

Tubule preparations were used for IHC. To obtain sperm samples, caudae epididymes were dissected and transferred to preheated (36-37°C) M2 medium, incised several times, squeezed with tweezers several times during a 30min incubation period at 36-37°C to ensure flushing out the whole sperm population including immotile sperm.

### Histology

Bouin-fixed testes and epididymes were paraffized, embedded in paraffin blocks and sectioned at 3 microns. Sections were deparaffinized and stained with the Periodic acid-Schiff (PAS) procedure. Stained sections were imaged under bright field at 20x and 63x magnification using a Leica DM5500 B microscope (Leica Microsystems, Wetzlar, Germany).

### Basic sperm analysis

Isolated spermatozoa were counted for 6-12 animals per genotype using a hemocytometer. Sperm motility was analyzed for 3-4 animals per genotype by taking 5-10, 3 second video clips per animal using a Basler acA1920-155ucMED camera (Basler AG, Ahrensburg, Germany). A minimum of 200 sperm per individual were analyzed and the percentage of motile sperm calculated. Sperm viability was analyzed by eosin-nigrosin staining for 3-4 animals per genotype. Approximately 1 x 10^6^ sperm were mixed with 50μl of eosin-nigrosin dye, incubated for 30 seconds, spread on slides, air-dried and cover-slipped. A minimum of 200 spermatozoa were analyzed under bright field (Leica DMIRB microscope) and the percentage of viable spermatozoa calculated.

### RNAseq and differential expression analysis

RNA was extracted from testes after removal of the tunica albuginea using the RNeasy kit (Qiagen, Hilden, Germany). RNA integrity (RIN) was determined using the RNA Nano 6000 Assay Kit with the Agilent Bioanalyzer 2100 system (Agilent Technologies, Santa Clara, CA, USA). RIN values ranged from 7.3–10 for all samples. RNA sample quality control and library preparation were performed by the University of Bonn Core facility for Next Generation Sequencing (NGS), using the QuantSeq 3’-mRNA Library Prep (Lexogen, Greenland, NH, USA). RNAseq was performed by the University of Bonn Core facility for Next Generation Sequencing (NGS) on the Illumina HiSeq 2500 V4 platform, producing >10 million, 50bp 3’-end reads per sample.

The samples were then mapped to the mouse genome (GRCm38.89) using HISAT2 2.1 (Kim et al. 2015). StringTie 1.3.3 (Pertea et al. 2015) was used for transcript quantification and annotation. Gene annotation was retrieved from the Ensembl FTP server (ftp://ftp.ensembl.org)(GRCm38.89). The python script (preDE.py) included in the StringTie package was used to prepare DEseq2-compatible gene-level count matrices for analysis of differential gene expression. Mapping to the *Prm2* genomic location was visualized using the Integrative Genomics Viewer (IGV; Robinson et al. 2011).

Differential expression (DE) was analyzed using DESeq2 1.16.1 (Love et al. 2014). The adjusted p-value (Benjamini-Hochberg method) cutoff for DE was set at < 0.05, log2 fold change of expression (LFC) cutoff was set at > 1. We performed GO term and pathway overrepresentation analyses on relevant lists of genes using the PANTHER gene list analysis tool with Fisher’s exact test and FDR correction (Mi et al. 2017).

### Sperm nuclear morphology

Nuclear morphology was analyzed for 3 individuals per genotype. Approximately 1.5 x 10^6^ sperm were fixed in methanol-acetic acid (3:1), spread onto a slide and stained with 4’,6-diamidino-2-phenylindole (DAPI) nuclear stain (ROTI^®^Mount FluorCare DAPI (Carl Roth GmBH, Karlruhe, Germany)). At least 200 stained sperm cells per individual were imaged at 100x magnification using a Leica DM5500 B fluorescent microscope. Nuclear morphology was analyzed using the stand-alone version of the Nuclear Morphology program by Skinner et al. (2019). The program allows for automated detection and morphological analysis of mouse sperm nuclei (among other species and cell types). It additionally provides options for clustering heterogenous populations by nuclear parameters and comparative analyses of nuclear morphology (Skinner et al. 2019). The parameters used for nucleus detection are shown in figure S3.

### Sperm basic nuclear protein extraction and analysis

Basic nuclear proteins were extracted described in Soler-Ventura et al. (2018). Briefly, approximately 10 x 10^6^ of swim-out sperm were washed in PBS and pelleted. The pellet was resuspended in buffer containing 1M Tris pH 8, 0.5M MgCl and 5ul Triton X-100. Subsequently the pellet was treated with 1mM PMSF in water inducing cell lysis. Treatment with EDTA, DTT and GuHCl induced DNA denaturation. Incubation at 37°C for 30min in presence of 0.8% vinylpyridine is necessary for mouse protamine separation on the subsequent protein gel. DNA is then precipitated by addition of EtOH and separated from the sample by centrifugation. Basic sperm nuclear proteins are then extracted and dissolved in 0.5M HCl, followed by protein precipitation with TCA, acetone washes and drying. The precipitated proteins are resuspended in sample buffer containing 5.5 M urea, 20% β-mercapto-ethanol and 5% acetic acid.

The samples were then run on a pre-electrophorized acid-urea polyacrylamide gel (AU-PAGE) (2.5 M urea, 0.9 M acetic acid, and 15% acrylamide/0.1% N,N-Methylene bisacrylamide, TEMED and APS). The extracted basic nuclear proteins migrate towards the negative pole at a 150V for 1h, 50min. The gels were stained with Coomassie Brilliant Blue (Sigma Aldrich, Taufkirchen, Germany) using standard procedures. The two main protamine bands can be observed in the bottom of the gel with mature-PRM2 corresponding to the upper and PRM1 the lower band (Ishibashi et al. 2010, Soler-Ventura et al. 2018). PRM2 precursor bands can be observed in the lower part of the gel above the mature-PRM2 band, if present (Yu et al. 2000, de Mateo et al. 2011). In the upper half of the gel, bands corresponding to other basic nuclear proteins, including histones can be found (see Soler-Ventura 2018). The densities of Coomassie stained bands were analyzed using ImageJ (1.52k, Schneider et al. 2012).

### Immunohistochemistry

PFA fixed testis and epididymis sections as well as EtOH fixed tubule preparations were used for immunofluorescent staining. Sections were deparaffinized in xylol and rehydrated. Sections and tubule preparations were washed in PBS and blocked for 30 min with normal horse serum (Vectorlabs, Burlingame, USA) at room temperature, followed by primary antibody incubation over night at 4°C. Antibodies and dilutions are shown in table S3. Slides were then double-stained with fluorescent secondary antibodies using the VectaFluor™ Duet Immunofluorescence Double Labeling Kit, DyLight^®^ 594 Anti-Rabbit (red), DyLight^®^ 488 Anti-Mouse (green) (Vectorlabs, Burlingame, USA), DAPI counterstained and coverslipped with ProLong™ Gold antifade reagent with DAPI (Thermo Fisher Scientific, Waltham, USA).

### Mass spectrometry and differential protein abundance analysis

Basic nuclear protein extractions were done for 3 individuals per genotype using approximately 10^6^ sperm. Extracted proteins were dissolved in sample buffer (5.5 M urea, 20% β-mercapto-ethanol and 5% acetic acid).

Peptide preparation: Protein solutions (5.5 M urea, 20% 2-mercaptoethanol, 5% acetic acid) were dried in a vacuum concentrator and subjected to in solution preparation of peptides. Proteins were dissolved in 50 mM acrylamide solution (Tris-HCl, pH=8) and alkylated for 30 min at RT. 1 μg of Trypsin were added for o/n proteolysis at 37°C. Dried peptides were dissolved in 10 μL 0.1% trifluoro acetic acid (TFA) and desalted with ZipTips (Waters GmbH, Eschborn, Germany) according to standard solid-phase extraction procedures. Equilibration and binding was done in presence of 0.1% TFA, washing with 0.1% formic acid (FA). Eluates (50% acetonitrile, 0.1% FA) were dried and stored at −20°C.

LC-MS measurements: Peptide separation was performed on a Dionex Ultimate 3000 RSLC nano HPLC system (Dionex GmbH, Idstein, Germany). The autosampler was operated in μl-pickup mode. Peptides were dissolved in 10 μl 0.1% FA (solvent A). 2 μL were injected onto a C18 analytical column (300 mm length, 75 μm inner diameter, ReproSil-Pur 120 C18-AQ, 1.9 μm). Peptides were separated during a linear gradient from 2% to 35% solvent B (90% acetonitrile, 0.1% FA) within 90 min at 300 nl/min. The nanoHPLC was coupled online to an Orbitrap Fusion Lumos mass spectrometer (Thermo Fisher Scientific, Bremen, Germany). Peptide ions between 300 and 1600 m/z were scanned in the Orbitrap detector every 3 seconds with R=120,000 (maximum fill time 50 ms, AGC target 400,000). Polysiloxane (445.12002 Da) was used for internal calibration (typical mass error ≤1.5 ppm). In a topspeed method peptides were subjected to higher energy collision induced dissociation (HCD: 1.0 Da isolation, threshold intensity 25,000, normalized energy 27%) and fragments analyzed in the Orbitrap with target 50,000 and maximum injection time 22 ms, R=15,000. Fragmented peptide ions were excluded from repeat analysis for 20 s.

Data analysis: Raw data processing and was performed with Proteome Discoverer software 2.5.0.400 (Thermo Fisher Scientific). Peptide identification was done with an in-house Mascot server version 2.6.1 (Matrix Science Ltd, London, UK). MS data were searched against *Mus musculus* sequences from SwissProt (2021/03, including isoforms), and contaminants (cRAP, Mellacheruvu et al. 2013). Precursor Ion m/z tolerance was 10 ppm, fragment ion tolerance 20 ppm. Tryptic peptides with up to two missed cleavages were searched. Propionamide on cysteines was set as static modification. Oxidation was allowed as dynamic modification of methionine, acetylation as modification of protein N-termini. Mascot results were evaluated by the percolator algorithm (Kall et al. 2008) version 3.05 as implemented in Proteome Discoverer. Spectra with identifications below 1% q-value were sent to a second round of database search with semitryptic enzyme specificity (one missed cleavage allowed). Protein N-terminal acetylation, methionine oxidation, carbamylation on lysine and N-termini were allowed as dynamic modifications. Actual FDR values were typically ≤0.5% (peptide spectrum matches), ≤1.0% (peptides), <1% (proteins). Proteins were accepted if at least two peptides with q-value <1% were identified. Summed abundances (areas of precursor extracted ion chromatograms of unique peptides) were used for relative quantification.

Differential abundance (DA) analysis: DA analysis was performed using the Bioconductor package proDA (Ahlmann-Eltze C 2021) using peptide spectrum matches (PSM) level data extracted from Protein Discoverer. Only proteins detected in all genotypes and all replicates with more than two peptides were included in the analysis. The data were log2 transformed and median normalized prior to DA analysis to ensure comparability. The proDA package is based on linear models and utilized Bayesian priors to increase power for differential abundance detection (Ahlmann-Eltze C 2021). Proteins with a log2 fold change (LFC) of >1 and false discovery rate adjusted p-value (FDR) <0.05 were considered differentially abundant compared to the WT. Plots were generated using the R-package ggplot2 (Wickam 2016).

### Generation of expression plasmids and transfection

*Prm2* and Prm2^Δc^ cDNA (GenBank: NM_008933.2) was amplified from C57Bl6 mouse testis cDNA using overhang primers introducing suitable restriction enzyme motifs (table S1) and cloned N-terminally in-frame with eGFP into the pEGFP-N3 vector (6080-1). Correct sequence and insertion were verified by sequencing.

Human Embryonic Kidney 293 (HEK293) cells were cultured in standard medium (DMEM, 10% FBS). HEK293 cells were transfected at 80% confluence with 3 μg of pPrm2-EGFP-N3 *(Prm2),* pPrm2^Δc^ -EGFP-N3 (Prm2^Δc^ sequence) or pEGFP-N3 with FuGENE^®^ HD Transfection Reagent (Promega, Madison, USA), according to the manufacturer’s instructions. At 12 hours post-transfection the medium was changed. Pictures were taken after 48 hours using a Leica DMIRB inverted microscope (Leica Microsystems, Wetzlar, Germany).

## Results

### Generation of gene-edited mouse lines

To delete cP2 in the reading frame of the PRM2 gene, we used CRISPR/Cas9 with templates catalyzing homology directed repair (HDR). In order to induce the deletion, we used two gRNAs targeting the 5’ and 3’ ends of the cP2 domain (Fig. 1A, Fig. S1). A single stranded DNA template encoding the 5’ and 3’ areas flanking the cP2 coding region enabling deletion of the cP2 coding region and an in-frame repair, generating an allele, where only mature PRM2 is expressed from the endogenous promoter, was added to the gene-editing reaction (Fig. S1). The sequence of the generated allele, named Prm2^Δc^, is shown in figure S1A and was registered with the mouse genomics database (MGI:6718282). Animals were generated, sequence validated and backcrossed to C57Bl/6J for at least 3 generations before analysis.

### Prm2^Δc/+^ as well as Prm2^Δc/-^ male mice are infertile

First, we subjected the Prm2^Δc/+^ mice to a fertility test. Five Prm2^Δc/+^ male mice were mated to ten WT females and the pregnancy/litters were recorded for five confirmed vaginal plugs per male (successful matings). We found that Prm2^Δc/+^ male mice were infertile, with no observed pregnancies in at least five confirmed matings each. This is in contrast to the deletion of the entire PRM2 gene, where Prm2^+/-^ male mice remained fertile (Schneider et al. 2016) (Fig. 1B). We hypothesized that an aberrant interaction between the newly generated mP2 and the PRM2 precursor expressed from the wildtype allele might lead to interference and be causative for infertility in these males. In order to test this, we bred Prm2^Δc/+^ females with Prm2^+/-^ males published by us (Prm2^Δ97bp^; Schneider et al. 2016, MGI:5760133) to generate Prm2^Δc/-^ mice. This results in male mice, in which only Prm2^Δc^ is present and expressed. However, Prm2^Δc/-^ males did not produce any litters in at least five confirmed matings each and can be considered infertile (Fig. 1B). This strongly suggests, that the cP2 domain is essential for murine spermiogenesis. If cP2 was non-essential these Prm2^Δc/-^ mice should be fertile, similar to Prm2^+/-^ males.

### mP2 is expressed in Prm2^Δc/-^ mice and transcriptional silencing does not seem to be disrupted

Since the Prm2^Δc/-^ mice allow for detection and validation of mP2 transcripts from the Prm2^Δc^ allele we, performed RNAseq on testis samples and analyzed expression of *Prm2.* By mapping the RNAseq reads to the *Prm2* genomic location, we were able to verify that the mP2 transcript was indeed expressed from the gene edited Prm2^Δc^ allele (Fig. S2). The expression levels of the *Prm2* transcripts were comparable in all genotypes (WT, Prm2^Δc/+^ and Prm2^Δc/-^), thus the alleles do not display a gene dosage effect (Fig. 1C).

In our previous study we found that in the Prm2^-/-^ testis, a much higher number of genes was differentially higher than lower expressed (81:13) indicating incomplete protamine-mediated transcriptional silencing (Schneider et al. 2020). Here, compared to WT, we found 36 genes differentially higher expressed and 42 genes differentially lower expressed for Prm2^Δc/+^ males and 26 genes differentially higher and 43 differentially lower for Prm2^Δc/-^ (Fig. 1D). This indicates that transcriptional silencing is not notably disrupted in males harboring the Prm2^Δc^ allele. No GO-term or pathway enrichment was found for analyzed gene sets. Lists of differentially expressed genes and statistics can be found in supplementary dataset S1.

### mP2 is detected in Prm2^Δc/+^ and Prm2^Δc/-^ spermatid nuclei, but is also found in cytoplasm and residual bodies

After confirming, that the gene-edited mP2-domain is expressed from the Prm2^Δc^ allele, we next addressed the question, whether mP2 can be detected in spermatids and is able to condense DNA. We therefore first performed IHC on PFA fixed testis sections. The epitope of the PRM2 antibody (Hup2B, Briarpatch Bio, Livermore, USA) is located in the first half of the mP2 domain and is able to detect both, the PRM2 protein generated from the wildtype and the gene-edited Prm2^Δc^ allele. As shown in figure 2A, PRM2 is detected in condensed spermatid nuclei of Prm2^Δc/+^ and Prm2^Δc/-^ males, similar to the wildtype. However, we additionally detected a signal in the cytoplasm of spermatids and residual bodies that is strongest in Prm2^Δc/-^ males and not found in wildtype (Fig. 2A).

**Figure 2.**
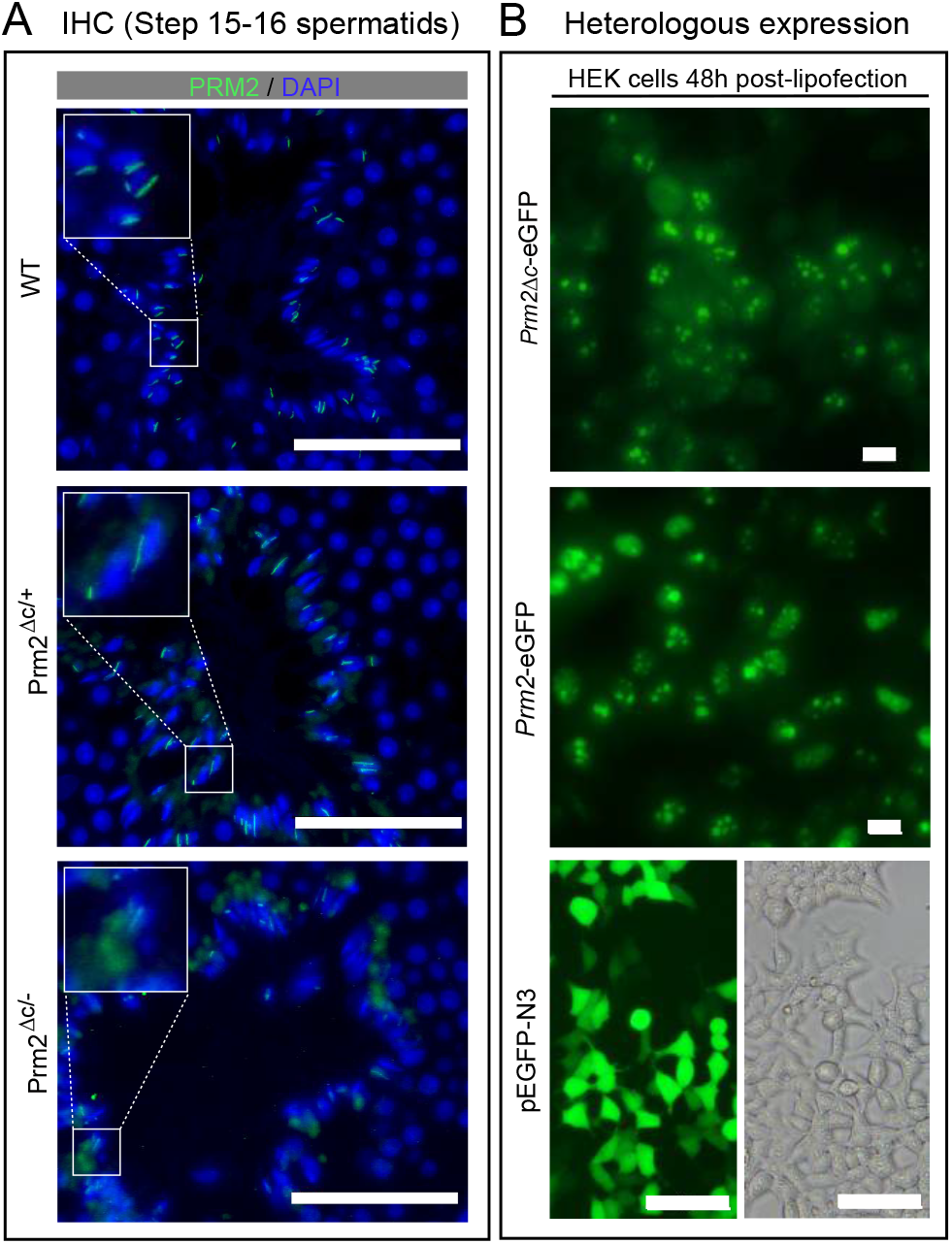
Localization and DNA condensing ability of mP2. A) Immunohistochemical fluorescent staining of PRM2 (WT, Prm2^Δc/+^) or mP2 (Prm2^Δc/+^, Prm2^Δc/-^) (green) in testis sections, counterstaining with DAPI (blue). Scale bar = 50μm. B) Heterologous expression of plasmids encoding eGFP tagged PRM2 (*Prm2*-eGFP) or mP2 (*mP2*-eGFP) in human embryonic kidney 293 (HEK) cells 48 hours post-transfection. Scale bar = 10μm.

To determine if mP2 is able to condense DNA, we expressed PRM2 and the mP2 sequence of the Prm2^Δc^ allele tagged with eGFP in HEK293 cells. The ability of PRM1 to bind to DNA and condense nuclei when expressed *in vitro* in somatic cells had been demonstrated before by Iuso et al. (2015) in sheep fibroblasts. Experiments revealed that PRM2-eGFP and Prm2^Δc^-eGFP locate to the HEK293 cell nuclei and are present after 48h as large speckles in the nucleus (Fig. 2B). We therefore concluded that mP2 is able to condense the nucleus of somatic cells *in vitro* and should therefore be able to contribute to spermatid chromatin condensation *in vivo.*

### Testis histology is inconspicuous, while mature sperm are inviable and immotile in Prm2^Δc/+^ and Prm2^Δc/-^ male mice

Next, we analyzed testis mass and examined histological sections to determine the nature of the infertility. Relative testes mass did not differ from the wildtype (Fig. 3A). Almost no viable and motile mature sperm were found in Prm2^Δc/+^ and Prm2^Δc/-^ males (Mean percent viable: Prm2^Δc/+^=1.7, SD=1.31; Prm2^Δc/-^=0; Mean percent motile: Prm2^Δc/+^=0.2, SD=0.45; Prm2^Δc/-^ =0.25, SD=0.5) (Fig. 3A, Fig. S4). Interestingly, sperm count is significantly reduced in Prm2^Δc/-^ but not in Prm2^Δc/+^ males compared to the wildtype (ANOVA: F(2)=10.87, p<0.001; Post-hoc Tukey HSD: WT vs. Prm2^Δc/+^: p=0.89, WT vs. Prm2^Δc/-^: p<0.001) (Fig. 3A). To determine if the loss of cP2 affects spermiogenesis, testis and epididymis histology was evaluated by PAS staining of Bouin-fixed sections. Testis histology was inconspicuous and spermatogenesis seemed not to be affected in either genotype (Fig. 3B). Epididymis histology however, shows larger round cells and vacuole-like structures, which are indicative of spermatid degradation. This was more pronounced in Prm2^Δc/-^ males compared to Prm2^Δc/+^ males (Fig. 3B).

**Figure 3.**
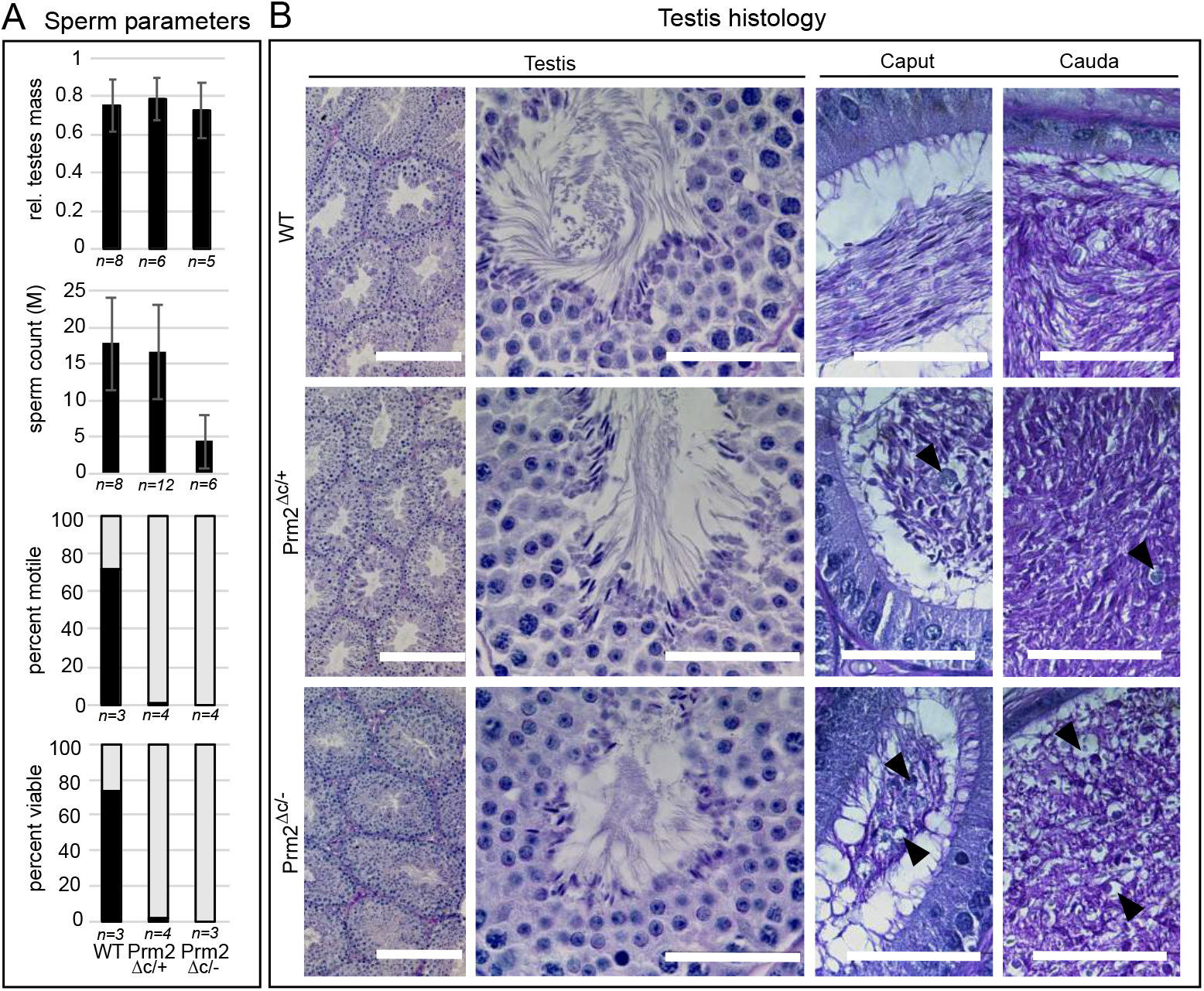
Sperm and testis parameters and histology. A) Barplots showing data for relative testes mass, mature sperm count, percentage of viable mature sperm (eosin-nigrosin assay) and percentage of motile mature sperm in Prm2^Δc/+^ and Prm2^Δc/-^ mice compared to wildtype. B) PAS staining of testis and epididymal sections of Prm2^Δc/+^, Prm2^Δc/-^ and WT males. Scale bar = 50μm (200μm for left column).

### Chromatin integrity is strongly affected and nuclear morphology is altered in Prm2^Δc/+^ and Prm2^Δc/-^ male mice

Since protamines condense sperm chromatin and were shown to influence sperm head morphology (Lüke et al. 2014a,b), mature sperm DNA integrity and nuclear morphology could be affected even though mP2 is able to condense DNA. We therefore extracted DNA from mature sperm and subjected it to agarose gel electrophoresis. DNA from sperm of Prm2^Δc/+^ males is completely fragmented (Fig. 4A). Schneider et al. (2020) was able to show that during epididymal transit, Prm2^-/-^ deficient sperm underwent ROS mediated destruction, leading to DNA and membrane degradation and immotility. We therefore stained sections of epididymides against 8-Oxo-2’-deoxyguanosine (8-OHdG), which indicates oxidative DNA damage. In the caput epididymis we detected only a slight increase of the 8-OHdG signal in Prm2^Δc/+^ and Prm2^Δc/-^ males compared to the wildtype. However, in the Prm2^Δc/+^cauda epididymis a strong increase in 8-OHdG was visible compared to the wildtype. The 8-OHdG signal in the Prm2^Δc/-^ cauda was less intense, which is likely due to the more severe degradation and reduced sperm count (Fig. 4B).

**Figure 4.**
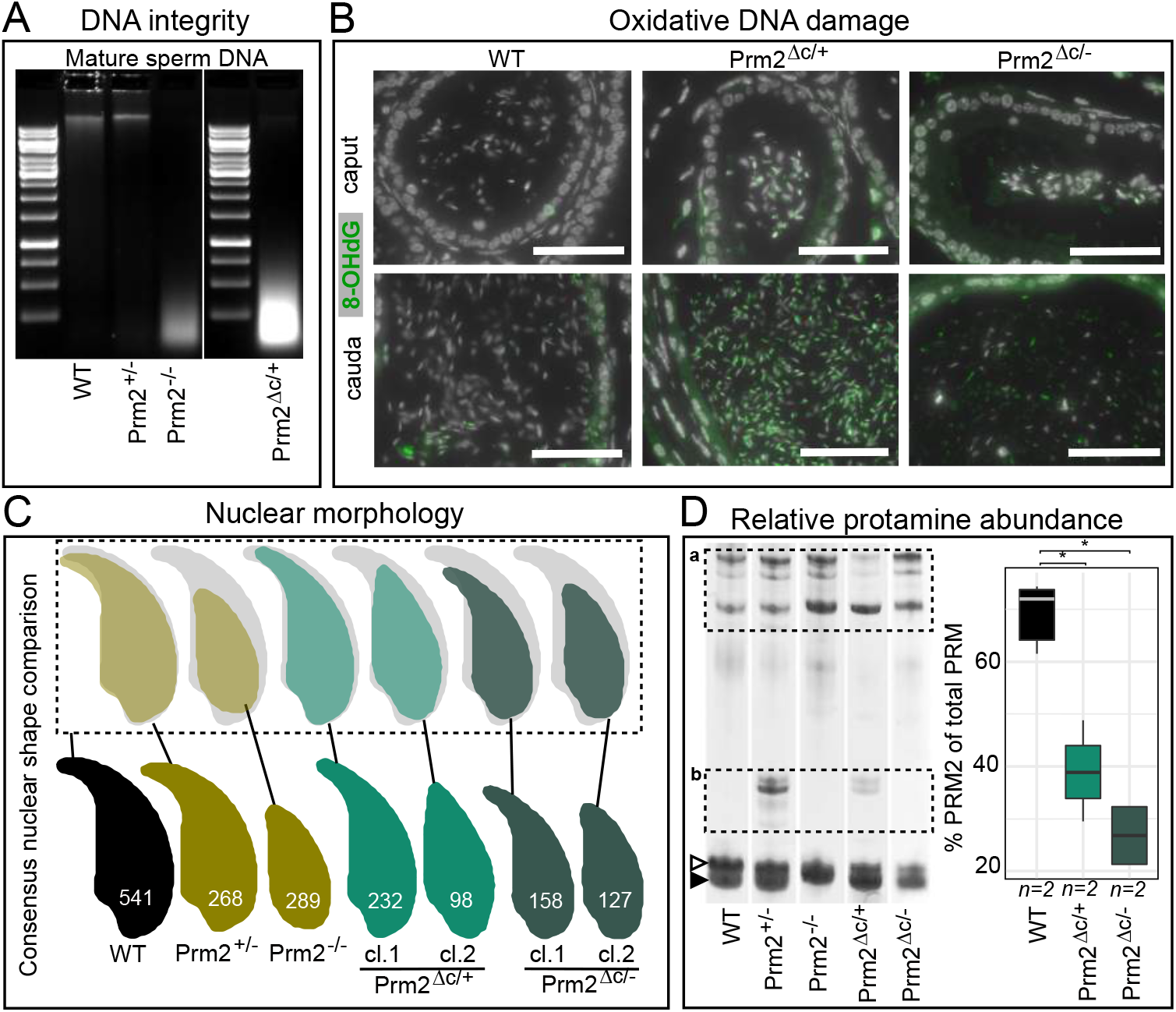
Chromatin integrity, nuclear morphology and protamine content. A) Agarose gel of DNA extracted from WT, Prm2^+/-^, Prm2^-/-^ and Prm2^Δc/+^ mature sperm. B) Immunohistochemical fluorescent staining of 8-Oxo-2’-deoxyguanosine (8-OHdG) (green) in caput epididymis (upper row) and cauda epididymis (lower row) of Prm2^Δc/+^, Prm2^Δc/-^ and WT. Counterstained with DAPI (pseudo-colored grey). Scale bar = 50μm. C) Comparison of mature sperm nucleus consensus shapes of WT, Prm2^+/-^, Prm2^-/-^, Prm2^Δc/+^ and Prm2^Δc/-^ resulting from nuclear morphology analysis. Numbers inside the consensus shapes indicate the number of nuclei assessed and assigned to the respective consensus shape cluster. Upper row shows the consensus shape of the different gene edited lines overlaid with the wildtype consensus shape. cl. = cluster. D) Representative lanes of acid-urea gel electrophoresis (AU-PAGE) of WT, Prm2^+/-^, Prm2^-/-^, Prm2^Δc/+^ and Prm2^Δc/-^ mature sperm basic nuclear protein extractions. a=non-protamine basic nuclear proteins, b=PRM2 precursors, open arrowhead indicates mature PRM2 band, solid arrowhead indicates PRM1 band. To the right: quantification of the percentage of PRM2 (including PRM2 precursors) of total protamine by band density analysis. Asterisk indicates significant difference.

Nuclear morphology analysis revealed aberrant nuclear morphology of mature sperm in Prm2^Δc/+^ and Prm2^Δc/-^ males. Sperm from both genotypes show a significantly reduced nuclear size compared to the WT (Table S2, Fig. 4C, Fig. S5). Prm2^Δc/+^ males show two clusters of nuclear shape, a slimmer nucleus with decreased hook curvature and a smaller hookless nucleus. This phenotype is even more severe in Prm2^Δc/-^ males (Table S2, Fig. 4C, Fig. S5). Of note, nuclear morphology of Prm2^Δc/-^ and Prm2^Δc/+^ deficient sperm also differs from Prm2^+/-^ and Prm2^-/-^ males.

The protamine ratio is flipped in Prm2^Δc/+^ and Prm2^Δc/-^ males Since the ratio between PRM1 and PRM2 is constant in mature sperm (in mice ~60% PRM2), and alterations of this ratio are associated with sperm defects and infertility (Corzett et al. 2002, Steger et al. 2008; García-Peiró et al. 2011), we next tested if male mice harbouring the Prm2^Δc^ allele display alterations of the PRM1/PRM2 ratio. Interestingly, acid-urea gel electrophoresis (AU-PAGE) of mature sperm basic nuclear proteins showed several bands corresponding to PRM2 precursors in Prm2^Δc/+^ males. This was also the case in Prm2^+/-^ males.

Comparing the densities of the PRM1 band relative to mP2 and PRM2 precursor bands, we found the PRM ratio to be strongly altered, showing a lower percentage of PRM2 (including precursors) compared to the wildtype in Prm2^Δc/+^ and Prm2^Δc/-^ males (Mean %PRM2: WT=69,29; Prm2^Δc/+^=42,07; Prm2^Δc/-^=26.78)(ANOVA: F(2)=18.98,p=0.009; Post-hoc Tukey HSD: WT vs. Prm2^Δc/+^: p=0.04, WT vs. Prm2^Δc/-^: p=0.008). This does not seem to be the case in Prm2^+/-^ males, for which we find the percentage of PRM2 (including precursors) to be similar to the wildtype (%PRM2=66.57 (n=1)) (Fig. 4D, Fig. S7). These data, together with the mP2 signal detected in condensing spermatid cytoplasm (Fig. 2A), strongly suggest, that loss of the cP2 domain leads to a reduction of PRM2 associated with DNA.

### Histone-to-protamine transition is incomplete in Prm2^Δc/+^ and Prm2^Δc/-^ males

Since we found the relative level of PRM2 to be reduced, and PRM2 aberrantly located in the cytoplasm and residual bodies of condensing/condensed spermatids in the testis, we next evaluated the histone-to-protamine transition. To this end, we performed IHC staining to detect histone H3, transition protein 1 (TNP1) and PRM1 in testis and epididymis sections. In Prm2^Δc/+^ male mice PRM1 staining intensity and localization is comparable to the WT. However, PRM1 staining seems to be lost in Prm2^Δc/-^ sperm from cauda epididymis (Fig. S6, Fig. 5). This is most likely due to the strong degradation of the sperm and its DNA and low sperm count in this genotype. We did not find any apparent increase in total histone H3 signal (Fig. S6, Fig. 5), However, in contrast to WT, TNP1 was retained in both Prm2^Δc/+^ and Prm2^Δc/-^ males in step 15-16 spermatids and caput epididymal sperm. In cauda epididymal sperm however, we did not find any visible signal of TNP1 (Fig. S6, Fig. 5). This indicates, that loss of cP2 leads to transition protein retention.

**Figure 5.**
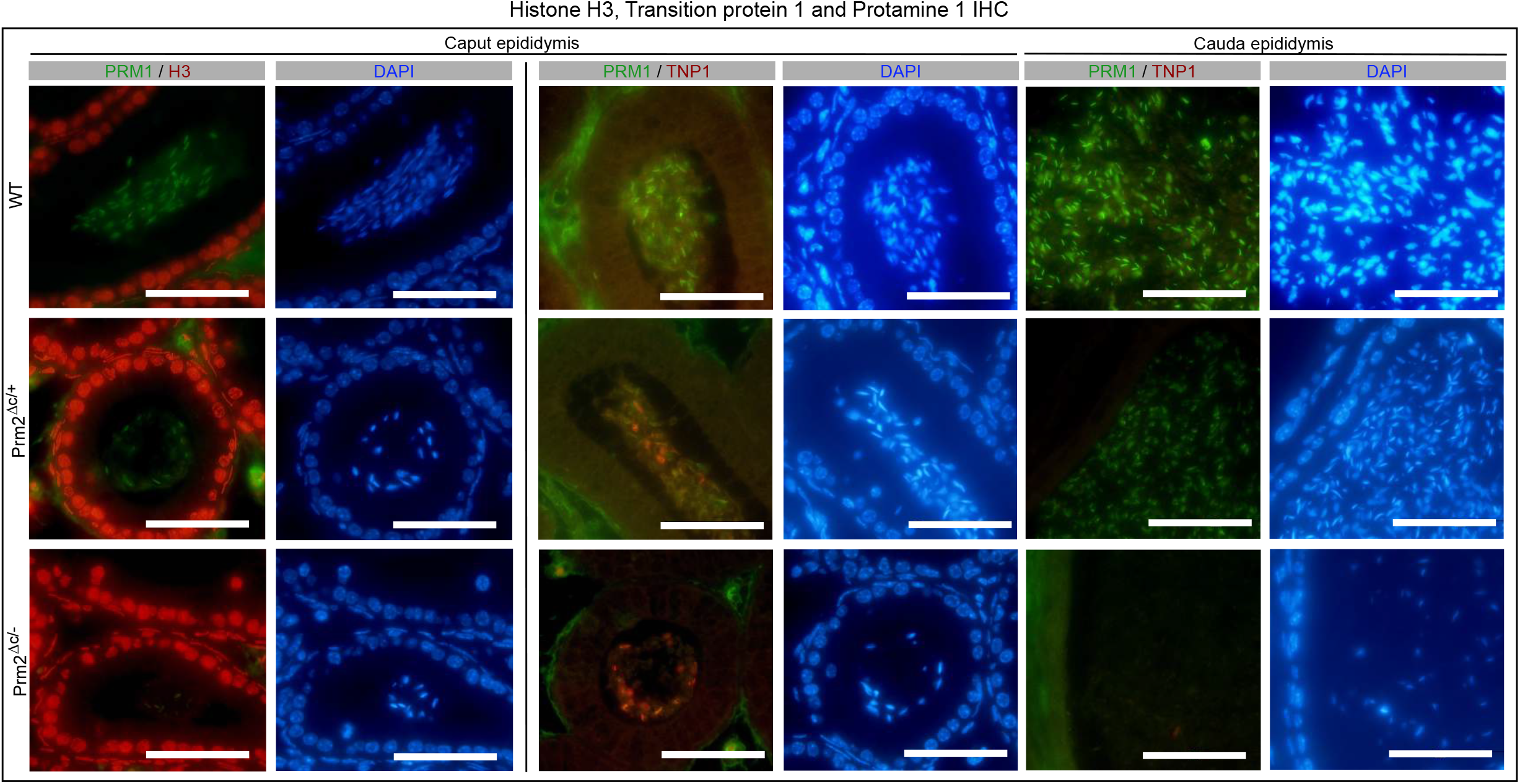
Histone H3, transition protein 1 and protamine 1 staining. Two left columns: Immunohistochemical fluorescent staining of Histone H3 (H3) (red) and protamine 1 (PRM1) (green) in WT, Prm2^Δ/+^ and Prm2^Δ/-^ caput epididymis. Counterstained with DAPI (blue). Scale bar = 50μm. Columns to the right: Immunohistochemical fluorescent staining of transition protein 1 (TNP1) (red) and protamine 1 (PRM1) (green) in WT, Prm2^Δ/+^ and Prm2^Δ/-^ caput and cauda epididymis. Counterstained with DAPI (blue). Scale bar = 50μm.

In order to further investigate histone retention and alteration in nuclear protein content we performed mass spectrometric analysis on mature sperm basic nuclear protein extracts and analyzed differential abundance (DA) of the detected proteins. Compared to wildtype, we found 14 proteins to be DA in Prm2^Δc/+^ sperm, 20 in Prm2^Δc/-^ males and 24 for Prm2^-/-^ sperm. Seven proteins were DA in all three comparisons, several of those associated with stress response and/or apoptosis (HSPA2, B2M, CLU) (Fig. 6A,B). Consistent with IHC results we did not find any histones that were significantly higher abundant in Prm2^Δc/+^ or Prm2^Δc/-^ males. We therefore conclude that histone retention is not increased in males harboring the Prm2^Δc^ allele. Interestingly in the Prm2^-/-^ samples we did find histones (H3f3, H3C, H4C) to be significantly higher abundant, indicating increased histone retention when PRM2 is completely lacking. Transition proteins were not detected. Lists of DA proteins including statistics can be found in supplementary dataset S2.

**Figure 6.**
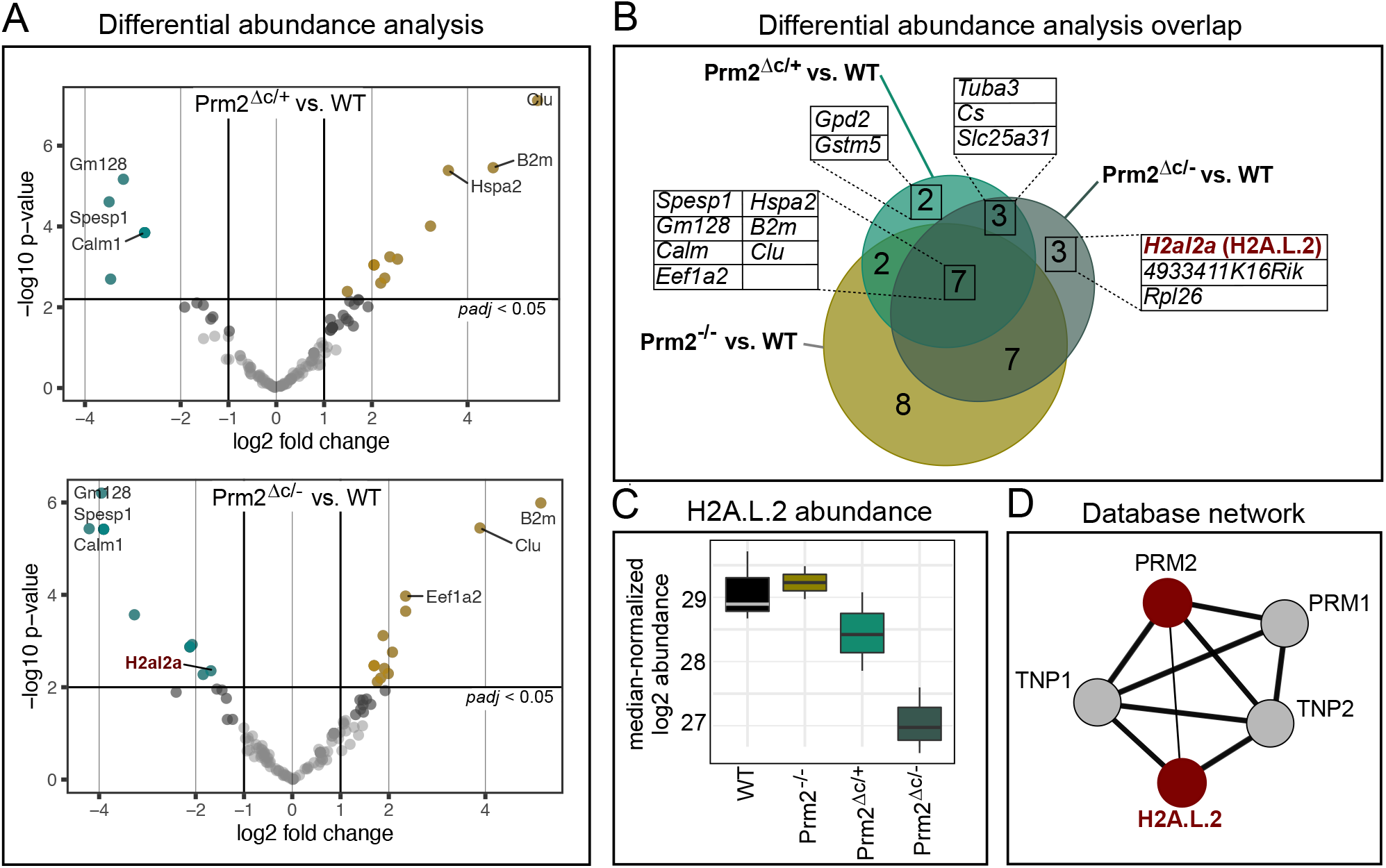
Differential abundance of mature sperm basic nuclear proteins. A) volcano plots showing differential abundance (DA) of basic nuclear proteins in Prm2^Δ/+^ compared to WT (upper plot) and Prm2^Δ/-^ compared to WT (lower plot). Significantly DA proteins are indicated in color (teal = lower abundant, yellow = higher abundant). Top DA proteins and proteins of interest are labeled with their corresponding gene symbol. B) Venn diagram showing the overlap between DA proteins found in the three different comparisons (WT vs. Prm2^Δ/+^, WT vs. Prm2^Δ/-^ and WT vs. Prm2^-/-^). Proteins present in overlaps of interest are listed with their corresponding gene symbol. H2A.L.2 is marked in red. C) Boxplot of median normalized log2 abundance of H2A.L.2 in WT, Prm2^-/-^, Prm2^Δ/+^ and Prm2^Δ/-^. D) Interaction network for H2A.L.2 extracted from String database (Szklarczyk et al. 2019).

Eight proteins were not DA in Prm2^-/-^ compared to WT, but in Prm2^Δc/+^ and/or Prm2^Δc/-^ males. Of these, RPL26 and GSTM5 are involved in DNA damage response and/or oxidative stress pathways, while TUBA3B and ANT4 *(Slc25a31)* are related to motility. Of note, citrate synthase (CS) is specifically higher abundant in Prm2^Δc/+^ and Prm2^Δc/-^ males. Most interestingly, we found the histone H2A variant H2A.L.2 to be significantly lower abundant in Prm2^Δc/-^ males, compared to the wildtype (Fig. 6A-D). H2A.L.2 is a spermatid/sperm-specific histone variant and a key player in histone-to-protamine transition. Together with TH2B it forms a nucleosome with an open chromatin structure allowing for loading of transition proteins followed by protamine recruitment and histone and transition protein eviction by protamines (Barral et al. 2017) (Fig. 6D). According to Hoghoughi et al. (2020) H2A.L.2 is retained in mature sperm in pericentric heterochromatin. We therefore investigated the presence and co-localization of H2A.L.2 and PRM2 in Prm2^Δc/-^ and Prm2^Δc/+^ males, by immunofluorescent staining of condensed step 15-16 spermatids from tubule preparations and mature sperm extracted from the cauda epididymis. In the wildtype, we found a moderately strong signal for H2A.L.2 in the pericentric region of the nucleus, with PRM2 signal in the whole nucleus (Fig. 7a-b,g-h). However, in Prm2^Δc/+^ and Prm2^Δc/-^ step 15-16 spermatids the H2A.L.2 signal is stronger compared to the WT and localized in DAPI-bright foci in the nucleus (Fig. 7c-f). DAPI-bright regions in nuclei usually correspond to heterochromatin. In mature sperm, however, the H2A.L.2 signal is almost completely lost in Prm2^Δc/+^ and Prm2^Δc/-^ males, consistent with the lower abundance found in mass spectrometric analysis. The PRM2 signal is diffuse and distributed along the whole mature sperm cell, likely due to severe membrane damage and degradation (Fig. 7i-l).

**Figure 7.**
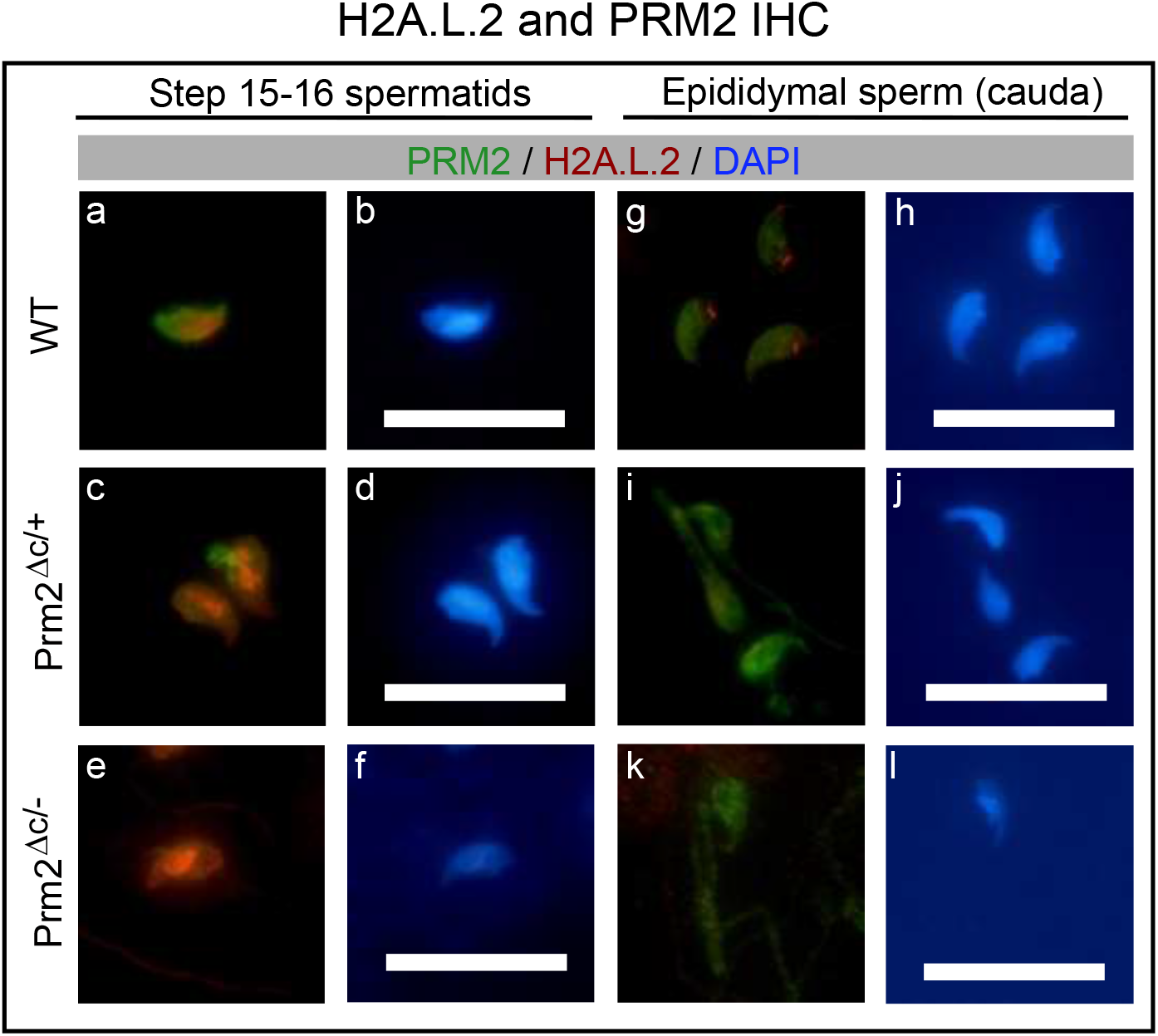
Immunohistochemical staining of H2A.L.2 and PRM2. Immunohistochemical fluorescent staining of H2A.L.2 (red) and PRM2 (green) in WT, Prm2^Δ/+^ and Prm2^Δ/-^ step 1516 spermatids from tubule preparations (two left columns) and mature sperm extracted from cauda epididymis (two right columns). Counterstained with DAPI (blue). Scale bar = 20μm.

## Discussion

The crucial role of protamines in the process of sperm chromatin reorganization is well known. However, why two protamines are needed in some species, while PRM1 seems to be sufficient in others is still unclear. The main difference between PRM1 and PRM2 is the highly conserved N-terminal cleaved PRM2 domain (cP2). Its function remains elusive to date. Using gene-editing we generated mice lacking the cleaved-*Prm2* (cP2) domain. We show that the cP2 domain is indispensable for PRM2 function and required for male mouse fertility. Mice heterozygous for the deletion of the domain display inviable and immotile sperm. Loss of cP2 leads to severe retention of transition proteins and reduced incorporation of PRM2 into nucleoprotamine. Instead, mature-PRM2 (mP2) is aberrantly found in spermatid cytoplasm and residual bodies. While overall histone retention is not increased, the histone variant H2A.L.2 is less abundant in mature sperm deficient in cP2.

Firstly, mP2 expressed from the Prm2^Δc^ allele was detected in the nucleus of condensed spermatids of cP2 deficient mice and was shown to be able to condense somatic cell DNA *in-vitro* similar to PRM2, or PRM1 (Iuso et al. 2015). We therefore conclude that the mP2 domain produced by gene editing seems to maintain its main function during spermiogenesis in cP2 deficient mouse lines. Male Prm2^Δc/+^ mice are infertile, show complete fragmentation of mature sperm DNA, loss of viability and immotility of mature sperm. This is in contrast to PRM2^+/-^ males, which remained fertile (Schneider et al. 2016). In order to test, whether the infertility of Prm2^Δc/+^ mice is due to an aberrant interaction between mP2 and the wildtype PRM2 precursor, we bred the Prm2^Δc^ allele with the Prm2^Δ97bp^ mouse line generated and analyzed by Schneider et al. (2016, 2020). However, mice expressing only mP2 (Prm2^Δc/-^) were also infertile showing an even more extreme phenotype than Prm2^Δc/+^ males. Since Prm2^+/-^ male mice are reported to be fertile, these data clearly indicate, that loss of cP2 leads to male infertility in mice. We speculate, that this holds also true for all other species which harbor a functional PRM2 gene.

Since mature sperm chromatin is completely fragmented in cP2 deficient males and mature sperm are inviable and immotile similar to Prm2^-/-^ sperm, we suspected that sperm might undergo epididymal degradation mediated by oxidative stress. Schneider et al. (2020) showed that loss of PRM2 seems to lead to reduced antioxidant capacity of sperm, initiating an oxidative stress-mediated destruction cascade during epididymal transit. Indeed, cP2 deficient mice showed a strong increase in oxidative DNA damage in the cauda epididymis. Prm2^Δc/-^ sperm seem to be degraded to an extent that many of the caudal sperm are completely disintegrated, leading to a significantly reduced sperm count. We propose that this is not a specific effect of cP2 or PRM2 deficiency but an inherent epididymal mechanism evoked by DNA damage or aberrant protamination (or even otherwise damaged sperm, e.g.: membrane defects, aberrant morphology or sperm surface proteome). How sperm damage might be sensed by epididymal cells and what, in fact, initiates this cascade is an interesting avenue for further investigation.

Given the timing of *Prm2* expression and processing, the primary effects of cP2 loss are likely to be found during the transition from histone-bound to protaminized DNA in the final stages of spermiogenesis. Barral et al. (2017) describe this transition taking place by assembly of a histone (TH2B and H2A.L.2) -transition protein (TNP1 and TNP2) interface followed by protamine recruitment and processing. Protamines themselves subsequently replace histones. This process is disrupted in cP2 deficient mice.

Firstly, in cP2 deficient mice mP2 is detected in the cytoplasm and residual bodies in addition to the signal found in the nucleus, indicating that mP2 is not completely incorporated into the condensing chromatin. In consequence, the protamine ratio changes from 2:1 in wildtype to 1:2-1:5 in cP2 deficient sperm. This indicates, that the cP2 domain facilitates incorporation of PRM2 and assembly of nucleoprotamine.

Secondly, we found retention of TNP1 in condensed testicular spermatids and caput epididymal sperm, indicating that the eviction of TNP1 is hampered due to the loss of cP2 or that TNP1 is binding DNA in a competitive manner. Of note, a recent study investigating a single residue mutation in PRM1 showed increased histone retention, but no disturbances in transition protein retention in such mice (Moritz et al. 2021). Transition proteins are believed to aid in chromatin condensation by stabilizing the DNA in a non-supercoiled state, cooperating with topoisomerases to relieve torsional stress and to be involved in DNA repair during chromatin condensation (Singh and Rao, 1988 Lèvesque et al. 1998, Akama et al. 1999). Loss of either transition protein results in incomplete PRM2 processing (Yu et al. 2000, Shirley et al. 2004, Zhao et al. 2004) and TNP2 was shown to interact with PRM2 (Barral et al. 2017). This demonstrates that transition proteins are required for proper PRM2 processing during nucleoprotamine assembly. The fact that we find severe retention of TNP1 in condensed spermatids indicates, that the cP2 domain in turn, is required for the proper processing (i.e. eviction) of TNP1.

Surprisingly, however, overall histone retention was not increased. This was confirmed by H3 immunostaining and mature sperm basic nuclear protein abundance analysis. Interestingly, we do find an increased retention of H3 and H4 variants in Prm2^-/-^ deficient sperm. The data from the H3 immunostainig indicate, that the global amount of retained histones seems unaffected, and hence not controlled by protamine ratio or PRM2 processing.

The retention of transition protein detected in cP2 deficient mice seems to go along with altered retention of H2A.L.2. During the first steps of chromatin condensation H2A.L.2, together with TH2B provides an open chromatin interface necessary for transition protein loading. Barral et al. (2017) showed, that deletion of H2A.L.2 leads to infertility, aberrant transition protein loading and disturbed processing of PRM2. In mature sperm H2A.L.2 was shown to be retained in pericentric heterochromatin (Govin et al. 2007, Hoghoughi et al. 2020). This retention seems to be lost in Prm2^Δc/-^ mature sperm, where H2A.L.2 was shown to be significantly lower abundant. However, a strong H2A.L.2 signal can be observed in Prm2^Δc/-^ condensed spermatids in the testis, were it overlaps with atypical speckles of bright DAPI signal. Thus, it seems that H2A.L.2-transition protein complexes are retained in aberrant clumps of heterochromatin in cP2 deficient mice.

Protamines have a strong electrostatic attraction to DNA due to arginine clusters (Moritz et al. 2021). This allows protamines to condense DNA even in the absence of spermatid specific histones and transition proteins, as shown in *in-vitro* assays in somatic cells and in this study (Iuso et al. 2015). However, uncontrolled or unbalanced binding of protamines might lead to strong torsional stress and DNA damage. A controlled stepwise chromatin condensation therefore could be required to maintain chromatin integrity. Moritz et al. (2021) recently showed that the PRM2 precursor has a lower DNA binding affinity than mP2, leading to faster DNA condensation by mP2. Barral et al. (2017) suggested that transition proteins buffer protamine incorporation by allowing for ordered protamine loading and (or possibly through) PRM2 processing. By losing the cP2 domain this processing step is skipped. We therefore propose, that the aberrant interaction with transition proteins due to cP2 deficiency leads to random mP2 binding, leading to strong hypercondensation of open chromatin resulting in DNA strand breaks. Transition protein-loaded chromatin, however does not allow for mP2 loading, leading to incomplete mP2 incorporation and retention of H2A.L.2 – transition protein complexes in aberrantly located heterochromatin foci, that are lost by DNA degradation during epididymal transit.

In conclusion, our results show that the cleaved domain of PRM2 is essential for sperm function and fertility. Loss of the domain leads to incomplete protamination, a switch in the protamine ratio and transition protein retention in epididymal sperm. During epididymal transit cP2 deficient sperm are degraded, seemingly by ROS mediated damage, leading to complete DNA fragmentation. cP2 seems to be necessary for correct interaction between the H2A.L.2 – transition protein complex and PRM2. We were able to provide a first glimpse into the function of cleaved PRM2 and PRM2 processing, that opens up multiple avenues for further investigation.

## Acknowledgments

This study was supported by grants from the Deutsche Forschungsgemeinschaft (DFG) to L.A. (AR 1221/1-1) and H.S. (Scho 503/15-2). L.A. was further supported by the University of Bonn Medical school supplementary research grant program *FEMHABIL.* We thank Gaby Beine, Angela Egert and Andrea Jäger for excellent technical assistance. Protein identifications were done at the Core Facility Mass Spectrometry, Institute of Biochemistry and Molecular Biology, Medical Faculty, University of Bonn. Mass spectrometer funded by the Deutsche Forschungsgemeinschaft (DFG) – Projektnummer 386936527. RNAseq was done at University of Bonn Core facility for Next Generation Sequencing (NGS). We thank Saadi Khochbin and Sophie Barral for the gift of the H2A.L.2 antibody and excellent advice.

## Author contributions

L.A. and H.S. conceived the study and designed the experiments. L.A. and S.S. generated gene-edited mice. L.A. and G.E.M. analysed gene-edited mice. F.E.O. and I.N. generated IHC stainings. L.A. and H.S. drafted the manuscript. All authors read and approved the final manuscript.

