## Supplementary material for "Loss of the cleaved-protamine 2 domain leads to incomplete histone-to-protamine exchange and infertility in mice": Fig. S1

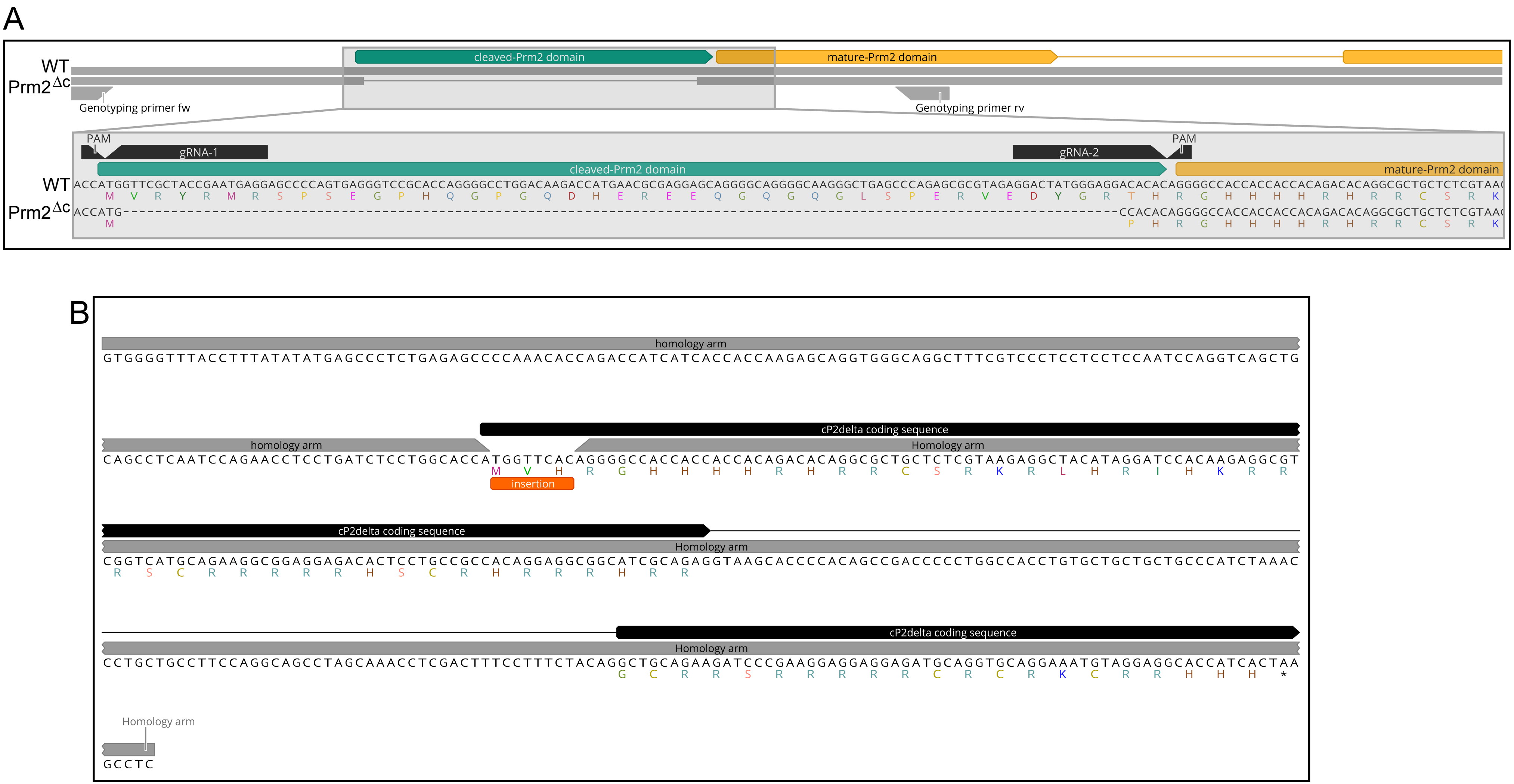


**Figure S1.** Gene editing. A) Schematic representation of CRISPR/Cas9 mediated gene editing. Gene-edited coding sequence and predicted translation are shown. B) ssODN repair template used during CIRSPR/Cas9 gene editing to evoke HDR mediates double strand break repair, fixing the startcodon and reading frame.
