## Supplementary figures and images for "Loss of the cleaved-protamine 2 domain leads to incomplete histone-to-protamine exchange and infertility in mice"

### Fig. S2

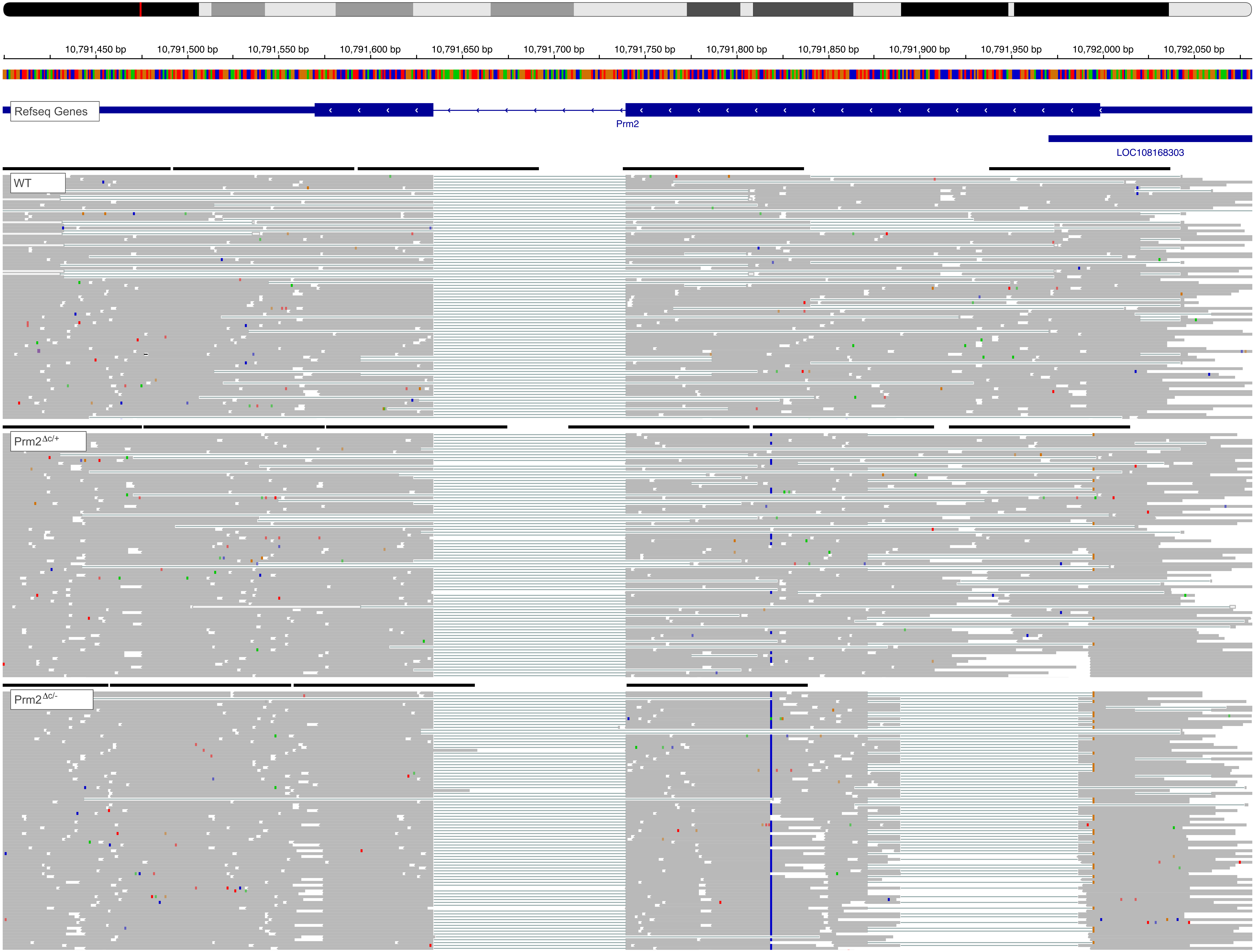


**Figure S2.** Representative cutout of mapping of RNAseq reads to the Prm2 genomic location.

### Fig. S4

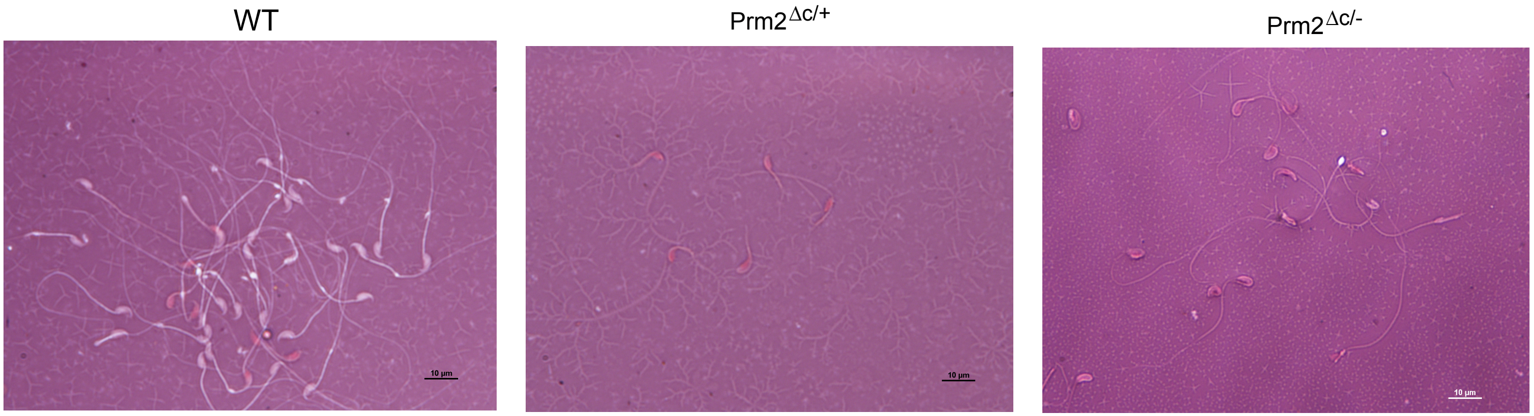


**Figure S4.** Representative pictures of eosin-nigrosin stained mature sperm

### Fig. S5

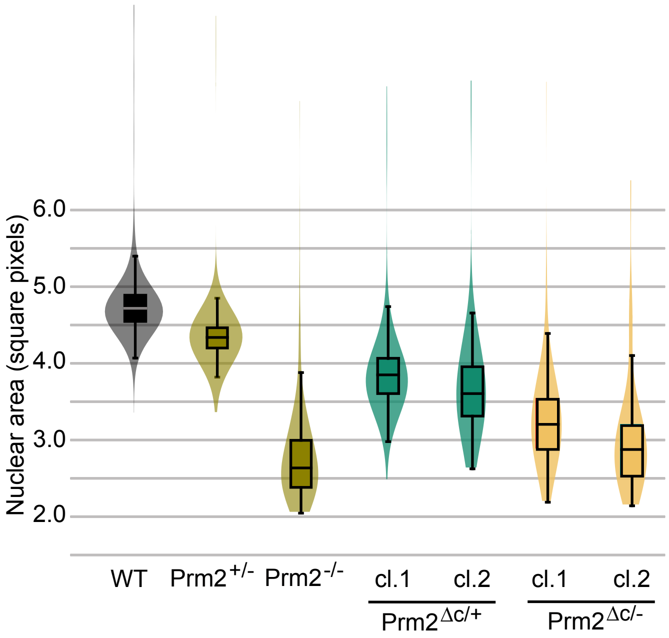


**Figure S5.** Boxplot of nuclear head area calculated during Nuclear Morphology Analysis
