## Supplementary material for "Loss of the cleaved-protamine 2 domain leads to incomplete histone-to-protamine exchange and infertility in mice": Fig. S3

| | |
|---|---|
| Image filtering | Kuwahara kernel: 7<br>Flattening threshold: 200 |
| Nucleus detection | Canny edge detection<br>Low threshold: 0.5<br>High threshold: 1.5<br>Kernel radius: 3.0<br>Kernel width: 16<br>Closing radius: 1 |
| Nucleus area | Min: 2000.0<br>Max: 10000.0 |
| Nucleus circularity | Min: 0.2<br>Max: 0.8 |

**Figure S3.** Detection parameters used for nuclus detection in the Nuclear Morphology program (Skinner et al. 2019)
