## Supplementary material for "Loss of the cleaved-protamine 2 domain leads to incomplete histone-to-protamine exchange and infertility in mice": Fig. S6

Histone H3, Transition protein 1 and Protamine 1 IHC

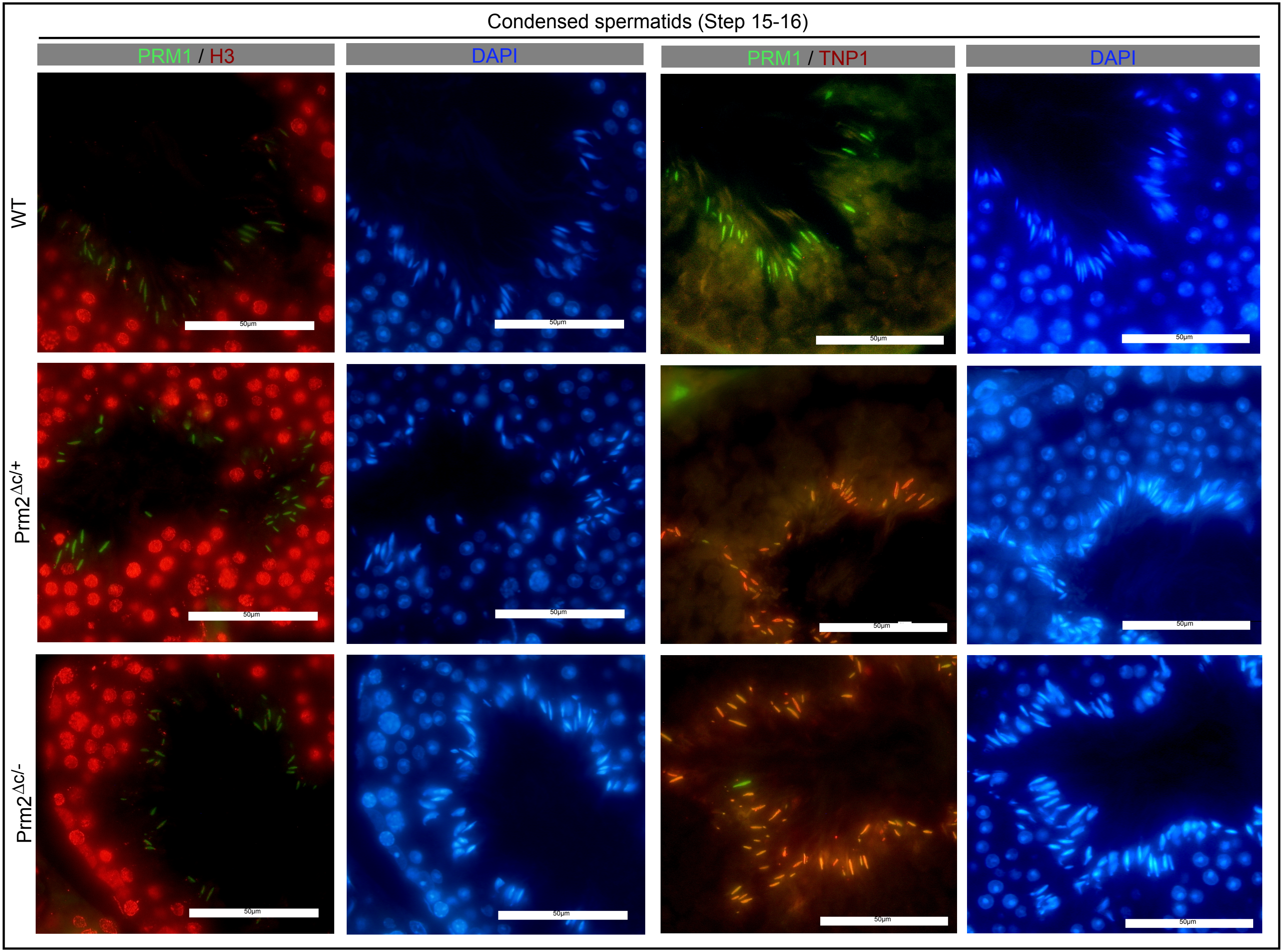


**Figure S6.** IHC of histone H3 (H3; red) and PRM1 (green) (two left columns) and TNP1 (red) and PRM1 (green) (two right columns) in testis sections showing step 15-16 spermatids in all three genotypes
