## Supplementary material for "Loss of the cleaved-protamine 2 domain leads to incomplete histone-to-protamine exchange and infertility in mice": Fig. S7

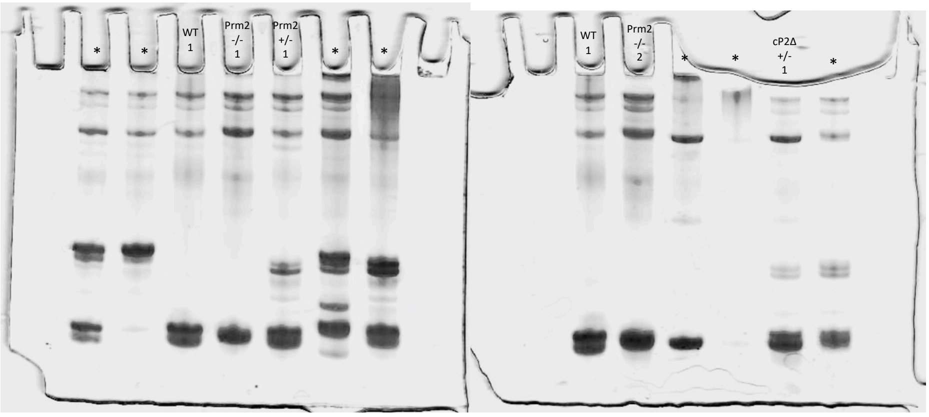


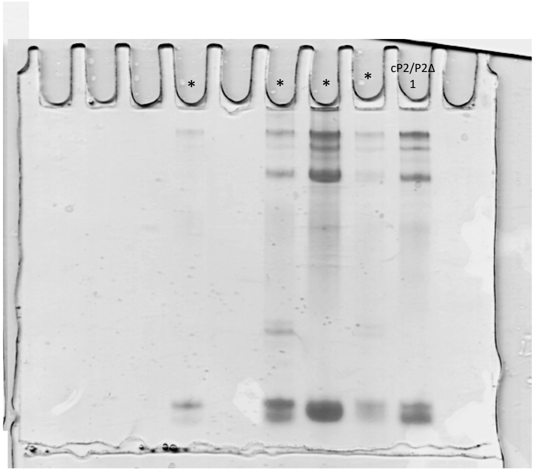


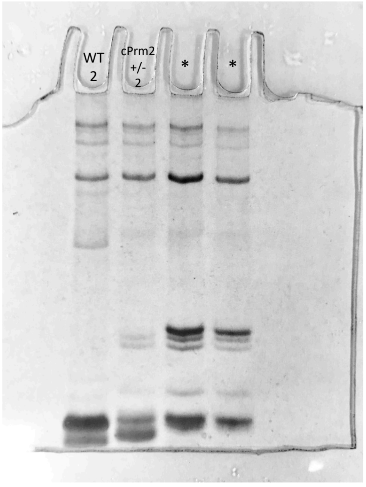


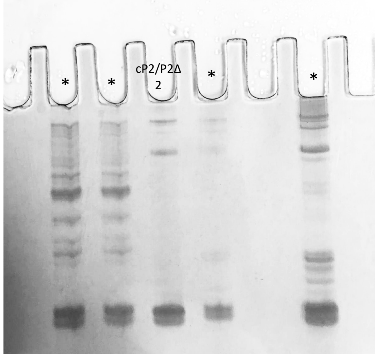


* Samples belonging to different experiments / not used in this study

**Figure S7.** Acid-Urea PAGE coomassie stained pictures. Lanes used for analysis of protamine ratio are indicated. Asterisk indicate lanes with samples belonging to different studies and not used in this study
